# The redefined role of PhaG and CoA ligases in medium-chain-length 3-hydroxy acid and polyhydroxyalkanoate production in *Pseudomonas putida*

**DOI:** 10.64898/2025.12.18.695248

**Authors:** A. R. Ola Pasternak, Walter T. Woodside, Eugene Kuatsjah, Sekgetho C. Mokwatlo, Jay D. Huenemann, William E. Michener, Stefan J. Haugen, Darren J. Parker, Alexis N. Williams, Kelsey J. Ramirez, Davinia Salvachúa, Gregg T. Beckham, Adam M. Guss

## Abstract

Under nutrient starvation conditions, many microorganisms such as *Pseudomonas putida* store excess carbon in medium-chain-length polyhydroxyalkanoate (*mcl*-PHA) granules. Biosynthesis of PHAs from the fatty acid biosynthetic pathway requires PhaG, which has long been thought to encode a 3-hydroxyacyl-ACP:CoA transferase, and PhaC, the PHA polymerase. Although this pathway has been extensively studied, the exact role of PhaG remains inconclusive. In this work we present *in vitro* biochemical and *in vivo* genetic evidence demonstrating that PhaG functions as a 3-hydroxyacyl-ACP thiolase, producing 3-hydroxyacids rather than 3-hydroxyacyl-CoA. We identified two CoA ligases, *fadD1* and *alkK*, essential for conversion of *mcl*-3-hydroxyacids to 3-hydroxyacyl-CoA, and thus PHA production. Taken together, this redefines the PHA biosynthetic pathway to include PhaG-dependent hydrolysis of hydroxyacyl-ACP, ligation of the 3-hydroxyacid to coenzyme-A, and polymerization by PhaC. Using these insights, we engineered *P. putida* to produce 3.7 g/L extracellular 3-hydroxyacids, which can be used for chemical synthesis of performance-advantaged polymers.

## Introduction

Microbial survival in variable environmental conditions has led to the evolution of myriad stress response strategies, with factors such as nutrient level variation and depletion often exhibiting strong regulatory effects on metabolism. Under conditions of nutrient (e.g., nitrogen or phosphorus) depletion and carbon excess, soil bacteria like *Pseudomonas putida* and *Cupriavidus necator* store excess carbon in granules of the intracellular polymer polyhydroxyalkanoate (PHA), which can later be utilized as a carbon source when nutrients are no longer limited.^1,2^ PHAs are classified as either short-chain-length (*scl*), typically composed of C4-C5 monomers or medium-chain-length (*mcl*), composed of C6-C14 monomers. One of the most well-studied *mcl*-PHA producers, *P. putida* KT2440, can produce *mcl*-PHAs from a variety of carbon sources, including from fatty acid ß-oxidation, and from sugars and aromatic feedstocks via *de novo* fatty acid biosynthesis.^2–5^

From fatty acid biosynthesis, the first step in PHA biosynthesis is thought to be transfer of a 3-hydroxyacyl group from acyl carrier protein (ACP) to coenzyme A (CoA) by PhaG, producing 3-hydroxyacyl-CoA **(Figure 1)**.^6^ Next, the 3-hydroxyacyl-CoA monomers are polymerized into PHA by the PHA polymerases, PhaC1 and PhaC2.^7^ PHAs can be hydrolyzed to 3-hydroxyacids (3-HA) by the PHA depolymerase, PhaZ.^1,8^ Despite this seemingly simple pathway, the catalytic function of PhaG remains unclear, due to differing reports in the literature. Previous *in vitro* studies suggest PhaG performs a 3-hydroxydecanoyl-ACP:CoA transferase function (3-hydroxyacyl-ACP + CoASH ↔ 3-hydroxyacyl-CoA + ACP).^6,9^ However, the PhaG transferase activity was only demonstrated in the reverse direction by monitoring the formation of 3-hydroxydecanoyl-ACP from 3-hydroxydecanoyl-CoA with *in vitro* assays^6^ and release of CoASH in kinetic studies^9^. More recently, *in vivo* studies using *Escherichia coli* strains engineered to produce PHAs via overexpression of *P. sp. 61-3 phaC1* and *P. putida phaG* led to extracellular accumulation of 3-hydroxydecanoic acid (3-HDA), consistent with thioesterase activity (3-hydroxyacyl-*Ec*ACP + H_2_O **→** 3-hydroxyacyl acid + *Ec*ACP), raising the possibility that PhaG is actually a thioesterase rather than a CoA transferase.^10,11^ Addition of a *P. putida* CoA ligase, *alkK* (PP_0763), resulted in increased PHA biosynthesis in *E. coli* further suggesting that AlkK, rather than PhaG, can perform CoA ligation, which is required for polymerization into PHA.^11^ Given the conflicting reports in the literature to date, identifying the definitive role of PhaG is critical to understand and harness the PHA biosynthetic pathway and will impact our understanding of how microbes store excess carbon and survive in environments with varying nutrient levels.

**Figure 1:**
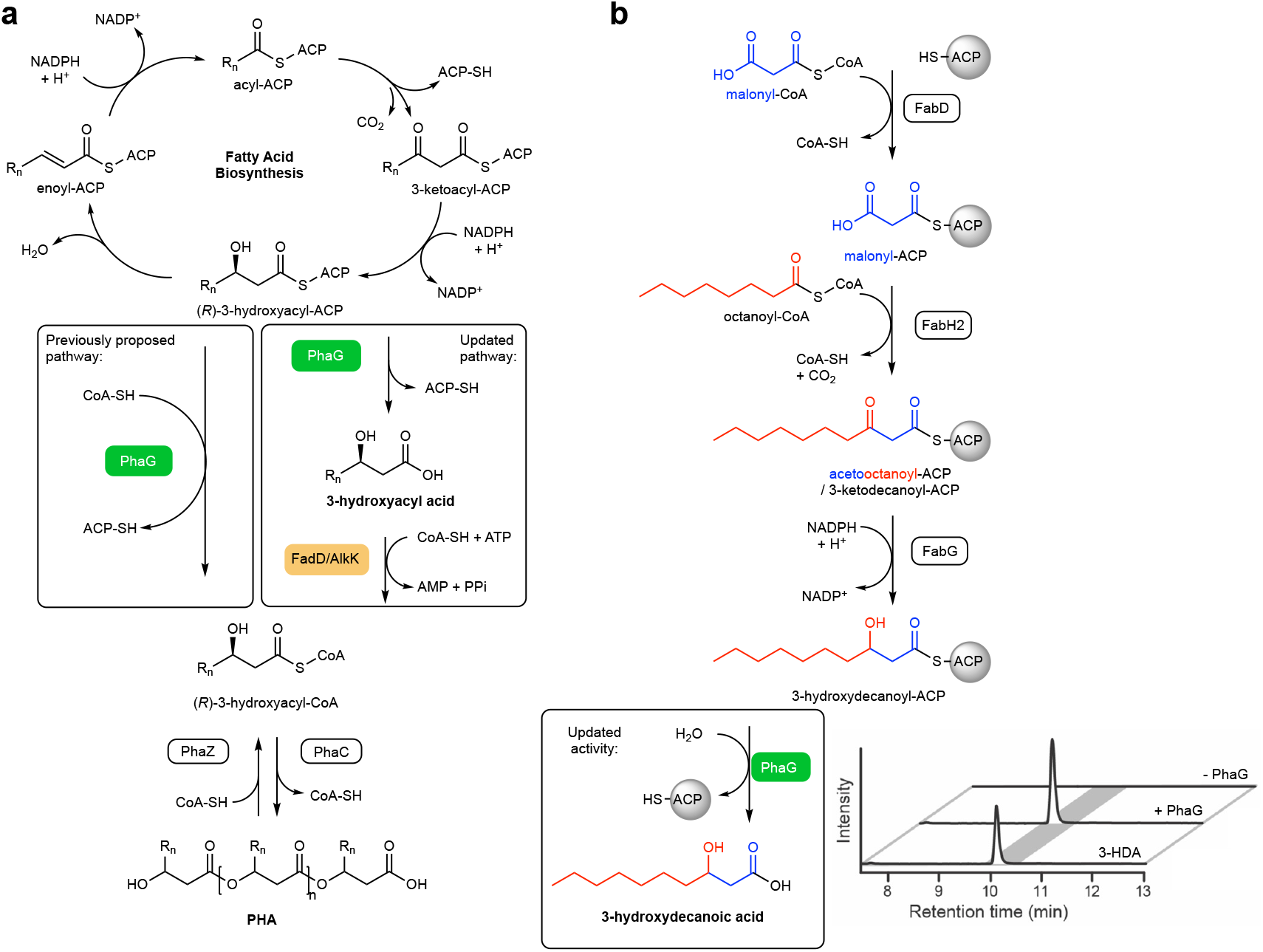
Pathways for production of 3-hydroxyacids and PHA. **(a)** *In vivo* pathway for PHA production in *P. putida*. The previously proposed pathway from fatty acid biosynthesis to 3-hydroxyacyl-CoA is in the left box, and an updated pathway with PhaG thioesterase activity is indicated in the right box. **(b)** The *in vitro* enzymatic cascade used to test PhaG activity. Bottom right, chromatographic resolution of the PhaG reaction. Traces: 3-HDA, 3-hydroxydecanoic acid standard; + PhaG, reaction containing *Ec*ACP, *Ec*FabD, malonyl-CoA, *Pp*FabH2, octanoyl-CoA, *Pp*FabG, NADPH, CoASH, and *Pp*PhaG; - PhaG, control reaction lacking *Pp*PhaG containing 3-hydroxydecanoyl-ACP and without PhaG.

Besides natural carbon storage products, PHAs have long been explored as alternatives to non-biodegradable, petroleum-based polymers, with applications in packaging, adhesives, and medical devices.^12–14^ PHAs are structurally diverse, biologically produced from various carbon sources including non-edible plant biomass,^3^ and in some forms, can biodegrade if leaked into the environment.^2,15,16^ Ultimately, a cost-effective PHA production platform could enable widespread use of these bio-based polymers. Biological production of PHA has been studied in many organisms. Notably *C. necator* is an effective producer of the *scl*-PHA, poly-3-hydroxybutyrate, while *mcl*-PHA production has been examined in *P. putida* and heterologously in *E. coli*, among other microbes.^5,11,17,18^ In addition to *in vivo* biological PHA production, designer PHAs can be chemically synthesized from 3-HA building blocks. Bioproduction of 3-HA monomers followed by chemical polymerization into PHAs may be an attractive route to bio-based polymers with enhanced properties. For example, PHA block copolymers can be designed with alternating stretches of different chain length *scl*-or *mcl*-PHA monomers, thus combining properties from both polymers to improve strength and flexibility.^18,19^ High titer biological production of *scl*-3-HA such as 3-hydroxybutyric acid (3-HB) has been achieved in *E. coli* from sodium gluconate and glucose.^20^ The engineered pathway condenses two acetyl-CoA units into 3-HB-CoA with a β-ketothiolase (*phbA*) and acetoacetyl CoA reductase (*phbB*) followed by a thioesterase (*tesB*) to release CoA and form 3-HB.^20^ However, achieving increased *mcl*-3-HA bioproduction titers has been challenging due to a lack of efficiently engineered metabolic pathways and has often relied upon *in vivo* production of *mcl*-PHA followed by either chemical or *in vivo* biological depolymerization.^21,22^ Overall, a detailed understanding of the PHA biosynthetic pathway could unlock the potential to optimize both 3-HA and PHA bioproduction, thus providing an alternative to traditional plastics manufacturing.

In this work, we re-examined the role of PhaG in PHA biosynthesis, first using biochemical assays, followed by systems biology and genetic manipulation. By discovering that PhaG functions as a thioesterase rather than a CoA transferase, we were able to rationally engineer *P. putida* to produce extracellular 3-HAs in bioreactor cultivations.

## Results

### PhaG hydrolyzes 3-hydroxydecanoyl-ACP to 3-hydroxydecanoic acid *in vitro*

We first sought to reproduce the proposed conversion of 3-hydroxydecanoyl-ACP to either 3-HDA or 3-hydroxydecanoyl-CoA *in vitro* to test whether PhaG functions as a thioesterase or acyl-transferase. Rather than measuring CoA liberation from the reverse reaction,^9^ a complete enzyme cascade reaction was performed using enzymes from *E. coli* (enzymes denoted with an *Ec* prefix) and *P. putida* (enzymes denoted with a *Pp* prefix) that were expressed and purified from *E. coli* to prime ACP with a 3-ketodecanoyl moiety, thus generating the physiological substrate for PhaG (**Figure 1b**).

To achieve this, holo-*Ec*ACP, *Ec*FabD, and *Pp*FabH2 were each purified to homogeneity. *Pp*PhaG, however, was only partially purified, as we were unable to replicate previously reported purification approaches.^23,24^ A small amount of soluble *Pp*PhaG was obtained when co-produced in conjunction with the chaperones GroEL and GroES. Further chromatographic purification of *Pp*PhaG reduced the protein yield, consistent with the inherent instability of the protein in the absence of the chaperones. Considering that *E. coli* does not produce PHA, we concluded that any observed PhaG-related activity in the assay is due to the contribution of the heterologous *Pp*PhaG.

The reaction was initiated by mixing *Ec*ACP, *Ec*FabD, and malonyl-CoA to produce malonyl-*Ec*ACP. Subsequently, *Pp*FabH2, a FabH homolog in *P. putida* that accepts *mcl*-acyl-CoA as an acyl donor, was added to catalyze a Claisen condensation between malonyl-*Ec*ACP and octanoyl-CoA, producing 3-ketodecanoyl-*Ec*ACP.^25^ Next, *Pp*FabG and NADPH were added to produce 3-hydroxydecanoyl-*Ec*ACP, followed by *Pp*PhaG addition. While we are unable to quantify 3-hydroxydecanoyl-CoA, in reactions containing *Pp*PhaG we detected 3-HDA produced at stoichiometric amounts to malonyl-CoA, octanoyl-CoA, and NADPH supplied in the reaction (1.1 ± 0.3 (mol/mol); *n* = 3), consistent with thioesterase activity (**Figure 1b**). No 3-HDA was detected in the control reactions without *Pp*PhaG.

### AlkK and FadD1 function as 3-hydroxyacyl acid-CoA ligases in *P. putida*

The *in vitro* data indicate that PhaG can hydrolyze 3-hydroxyacyl-ACP to free 3-HA. If this activity is also relevant *in vivo*, the resulting free 3-HA would need to be ligated to CoA to allow polymerization to PHAs. Because CoA ligation is also the first biochemical step in growth on fatty acids, we therefore hypothesized that identification of the CoA ligases needed for growth on 3-HA would also reveal the enzymes needed for PHA biosynthesis. We sought to identify these putative CoA ligases by looking at differential gene expression during growth on 3-HDA compared to acetate (**Figure 2a, Table S1**). *P. putida* encodes thirteen putative fatty acid CoA-ligases, of which nine are predicted to function on medium and long chain fatty acids, based on amino acid sequence similarity and annotations in both the BioCyc^26^ and the *Pseudomonas* Genome Database.^27^ Of these, only *fadD1* (*PP_4549*) was upregulated more than 2-fold during growth on 3-HDA compared to growth on acetate (10-fold increase), and *alkK* (*PP_0763*) was second highest with a 1.7-fold increase.

**Figure 2:**
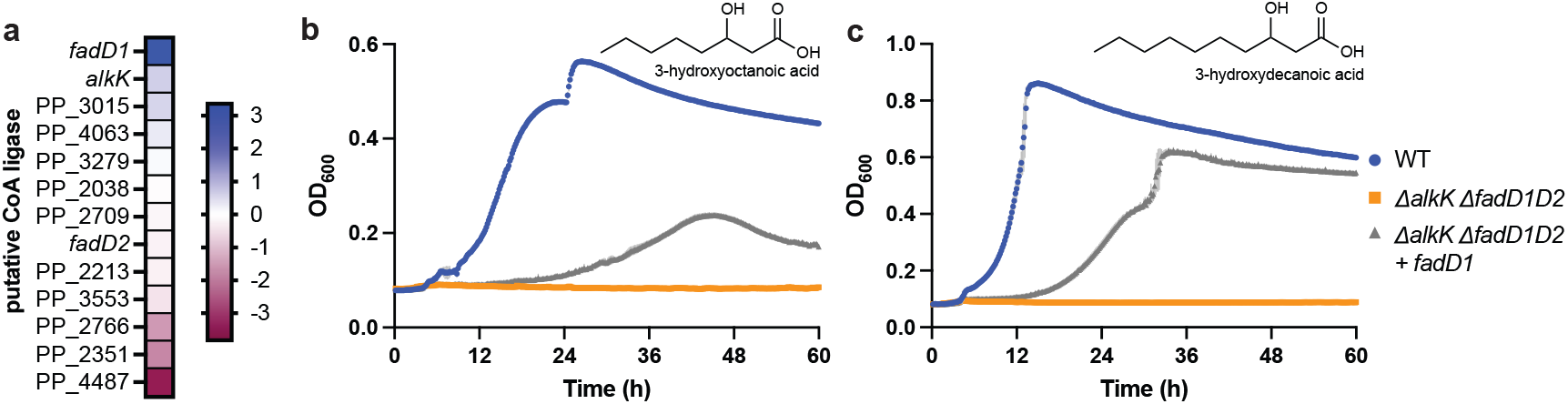
CoA ligases required for growth on *mcl*-3-HA. **(a)** Heat map (left) showing log_2_ change in transcript abundance of putative CoA ligases in WT *P. putida* when grown on 3-HDA relative to acetate (scale on right). Log_2_ values are shown in **Table S1**. Growth curves of *P. putida* mutant strains in modified M9 medium with the following 3-HA as the sole carbon source: **(b)** 3-HOA and **(c)** 3-HDA. Wild-type (WT) *P. putida* (blue), Δ*alkK*Δ*fadD1D2* (orange), and Δ*alkK*Δ*fadD1D2* + *fadD1* (grey). Results show the average of biological triplicates and error bars represent standard deviation. Source data are provided as a Source Data file.

To test the importance of these CoA-ligases for growth of *P. putida* on 3-HA, we deleted the gene cluster containing *fadD1* and the adjacent gene *fadD2* (*PP_4550*) with and without additional deletion of *alkK*, since *alkK* expression enabled PHA biosynthesis in *E. coli*.^11^ These strains were cultivated in medium containing 3-HDA or 3-hydroxyoctanoate (3-HOA) as the sole carbon and energy source. As a control, to verify that PhaG does not catalyze CoA-ligation *in vivo*, a Δ*phaG* strain was also created. The wild type and *ΔphaG* strains grew similarly well on 3-HDA, indicating CoA-ligation is still occurring without PhaG (**Figure S1b**). The strain with a single deletion of *alkK* also grew well, but the Δ*fadD1D2* strain grew substantially slower on 3-HDA and reached a lower final OD_600_ (**Figure S1a**), suggesting that *fadD1* is the major CoA ligase during growth on this *mcl*-3-HA, but that another CoA ligase exists that can serve the same function. Combining these two deletions (Δ*alkK*Δ*fadD1D2*) resulted in a strain completely incapable of growth on both 3-HDA and 3-HOA, with growth restoration upon complementation with a secondary copy of *fadD1* (**Figure 2b,c**). Our observed growth defects associated with *alkK* and *fadD1* deletion are consistent with previously reported fitness defects in *alkK* and *fadD1* mutant strains when grown on *mcl*-fatty acids, suggesting that AlkK and FadD1 also catalyze CoA ligation of C6 to C18 fatty acids.^28,29^ To confirm that CoA ligation is not impacted by enzymes downstream in the PHA pathway, *phaC1ZC2*, were deleted, and the resulting strain also grew on 3-HDA (**Figure S1b**). Together, this confirms that *fadD1* is critical for CoA ligation to 3-HAs.

### CoA ligase deletion mutants do not synthesize PHA

Based on the observations above, we further hypothesized that strains incapable of growth on 3-HAs would also not be capable of synthesizing PHAs because CoA ligation of 3-HA is required for polymerization into PHA by PhaC. To test this, we grew wild-type *P. putida*, Δ*alkK*Δ*fadD1D2*, and the *fadD1* complemented mutant with *p*-coumarate under nitrogen-excess and nitrogen-limited conditions, the latter which induces *mcl*-PHA production in the wild-type strain.^2^ First, we examined the cells by cryo-electron microscopy to look for PHA granules (**Figure S2**). As hypothesized, none of the strains exhibited substantial intracellular accumulation of PHA granules during growth with excess nitrogen, with the wild-type strain containing potentially small granules. Under N-limited growth, PHA granules were observed in the parent strain and the *fadD1*-complemented strain, but not in the Δ*alkK*Δ*fadD1D2* mutant, suggesting that this mutant does not make PHA.

To quantify the impact on PHA production, we measured PHA titer during growth on nitrogen-limited medium (**Figure 3a**). As expected, the wild-type strain produced 300.3 ± 12.4 mg/L PHAs, with 3-hydroxydecanoyl (C10) and 3-hydroxyoctanoyl (C8) monomers as the major components. Strains with mutations known to prevent PHA biosynthesis, Δ*phaG* and Δ*phaC1,Z,C2*, produced very low titers, 5.81 ± 0.35 mg/L and 3.69 ± 0.39 mg/L PHA, respectively, in nitrogen-limited medium. In the Δ*alkK*Δ*fadD1D2* strain, PHA titers were also very low (3.08 ± 0.095 mg/L). Upon complementation of *fadD1*, PHA production was largely restored (204.2 ± 4.11 mg/L) (**Figure 3a**), which further demonstrates that CoA ligation is required for PHA accumulation.

**Figure 3.**
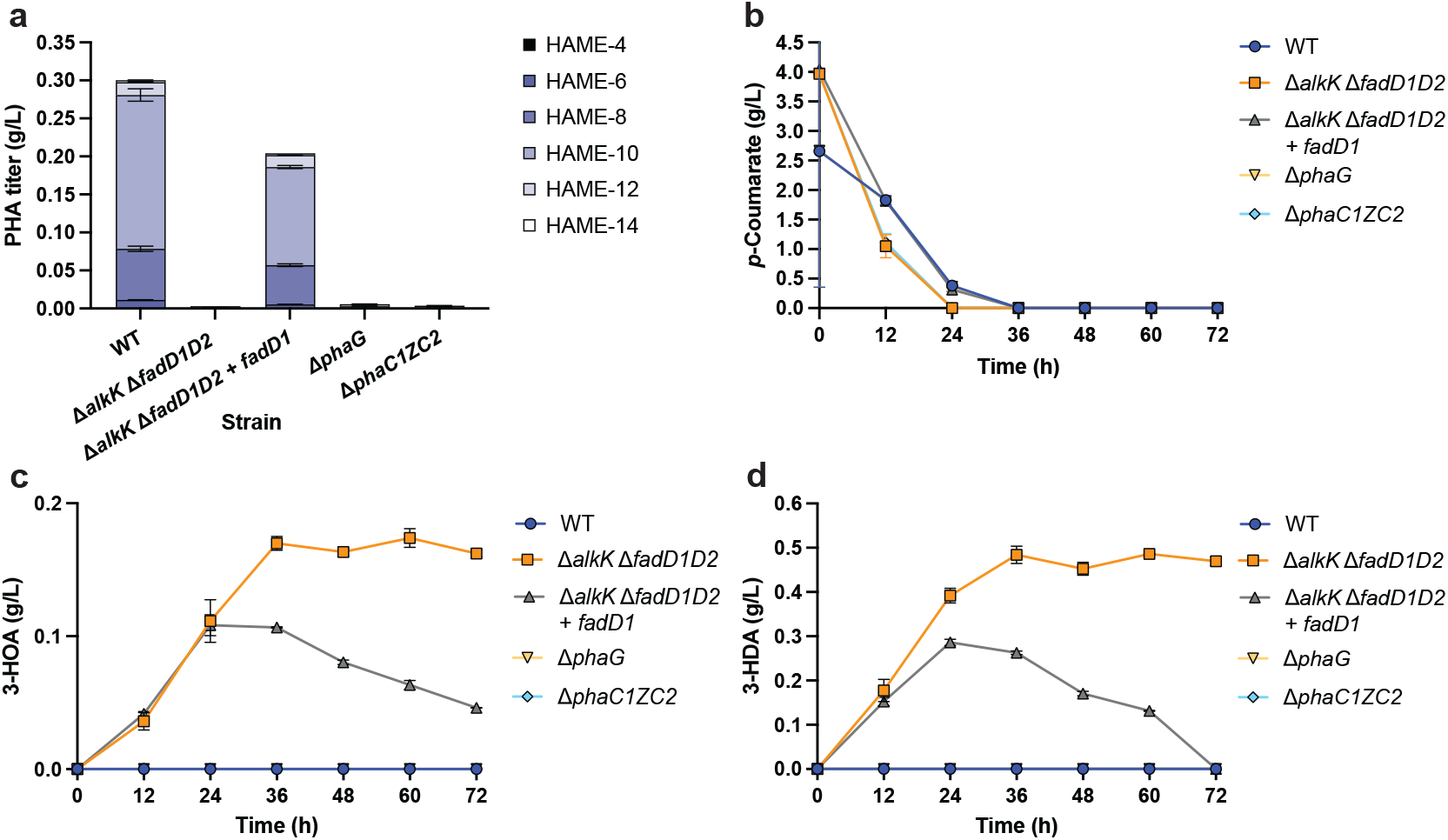
Production of PHA and 3-HA from *p*-coumarate in *P. putida* mutant strains. **(a)** PHA titers and composition, measured by hydroxyacyl methyl ester (HAME) derivatization of depolymerized PHA, **(b)** *p*-coumarate consumption by *P. putida* strains, **(c)** 3-HOA production titers, and **(d)** 3-HDA production titers. Data points are the average of triplicate cultures, and the error bars represent standard deviation. Source data are provided as a Source Data file.

### 3-HAs are secreted in CoA ligase mutants

After discovering that CoA ligase deletion mutants do not accumulate PHA, we sought to determine if these strains secrete PHA precursor products such as 3-HA instead. Following growth on excess *p*-coumarate and nitrogen-limited medium in shake flasks, HPLC analysis revealed no 3-HA production in cultures of the wild-type *P. putida* parent strain or the Δ*phaG* strain (**Figure 3c-d**), whereas the Δ*alkK*Δ*fadD1D2* strain accumulated 160 mg/L 3-HOA and 470 mg/L 3-HDA in the supernatant over 72 hours (**Figure 3c-d**). Complementation of *fadD1* into the Δ*alkK*Δ*fadD1D2* strain, generated a strain that transiently accumulated but then consumed 3-HA, further demonstrating that CoA-ligation of 3-HA is performed by FadD1 (**Figure 3c-d**).

We also considered an alternative hypothetical pathway in which PhaG is a CoA transferase and 3-HAs are produced via PhaZ-catalyzed PHA hydrolysis (**Figure S3**). In this potential pathway, CoA ligase deletion mutants would accumulate 3-HA because FadD1 does not reincorporate them after PHA hydrolysis. If this were correct, deletion of the PHA depolymerase (*phaZ*) in a Δ*alkK*Δ*fadD1D2* strain would eliminate 3-HA accumulation. However, the Δ*alkK*Δ*fadD1D2* Δ*phaZ* supernatant showed 3-HA levels similar to Δ*alkK*Δ*fadD1D2* (**Figure S4**), indicating PhaZ does not play a role in 3-HA production and further supporting our model that PhaG is a thiolase.

### Production of *mcl*-3-HA in bioreactors

Following substantial 3-HA production in shake flasks, we evaluated the capacity of *P. putida* Δ*alkK*Δ*fadD1D2* to produce 3-HA in bioreactors. A dissolved oxygen (DO)-stat fed-batch strategy was used to feed glucose in pulses to minimize 2-ketogluconate accumulation.^30^ By varying glucose and (NH_4_)_2_SO_4_ concentrations, three feeding media compositions with different carbon-to-nitrogen (C:N) molar ratios (20:1, 50:1, and a no-nitrogen case) were tested. The batch phase was initiated at a glucose concentration of 10 g/L, which was depleted after 9.5 h, triggering (DO)-stat controlled feeding. Cell growth in the batch phase reached a value of OD_600_ of ∼9 in all cases (**Figure 4a**), whereas cell growth varied significantly during the fed-batch phase. As expected, the C:N ratio of 20 led to the highest final OD_600_ (17.2 instead of 7.95 and 3.55 at C:N ratios of 50 and the no-nitrogen case, respectively) (**Figure 4a**). C:N ratios of 20 and 50 showed similar production profiles of total 3-HA, achieving a maximum titer of 2.23 g/L (**Figure 4b**). However, removing nitrogen altogether was detrimental, reaching a 3-HA titer of only 1.67 g/L (**Figure 4b**). In all three conditions, 3-HA production continues to increase after growth ceases, indicating production is not coupled to growth. Additionally, 3-HA composition primarily consisted of the C10 product, 3-HDA, with over 80% abundance (on a mass basis) for media conditions with C:N ratios of 50 and 20 and 68% when no nitrogen was fed (**Figure 4c**). The C8 product, 3-HOA was the second most abundant and the C6 3-HA was only produced at 3% for the condition where no nitrogen was present. Despite the absence of measurable glucose content in the bioreactor during the fed-batch phase (**Figure S6a**) 2-ketogluconate significantly accumulated in the bioreactors when no nitrogen was fed (up to 60.7 g/L at 49 h, almost the same molar concentration as glucose added) and at a C:N ratio of 50 (**Figure S6c**) (up to 12.9 g/L at 49 h).

**Figure 4.**
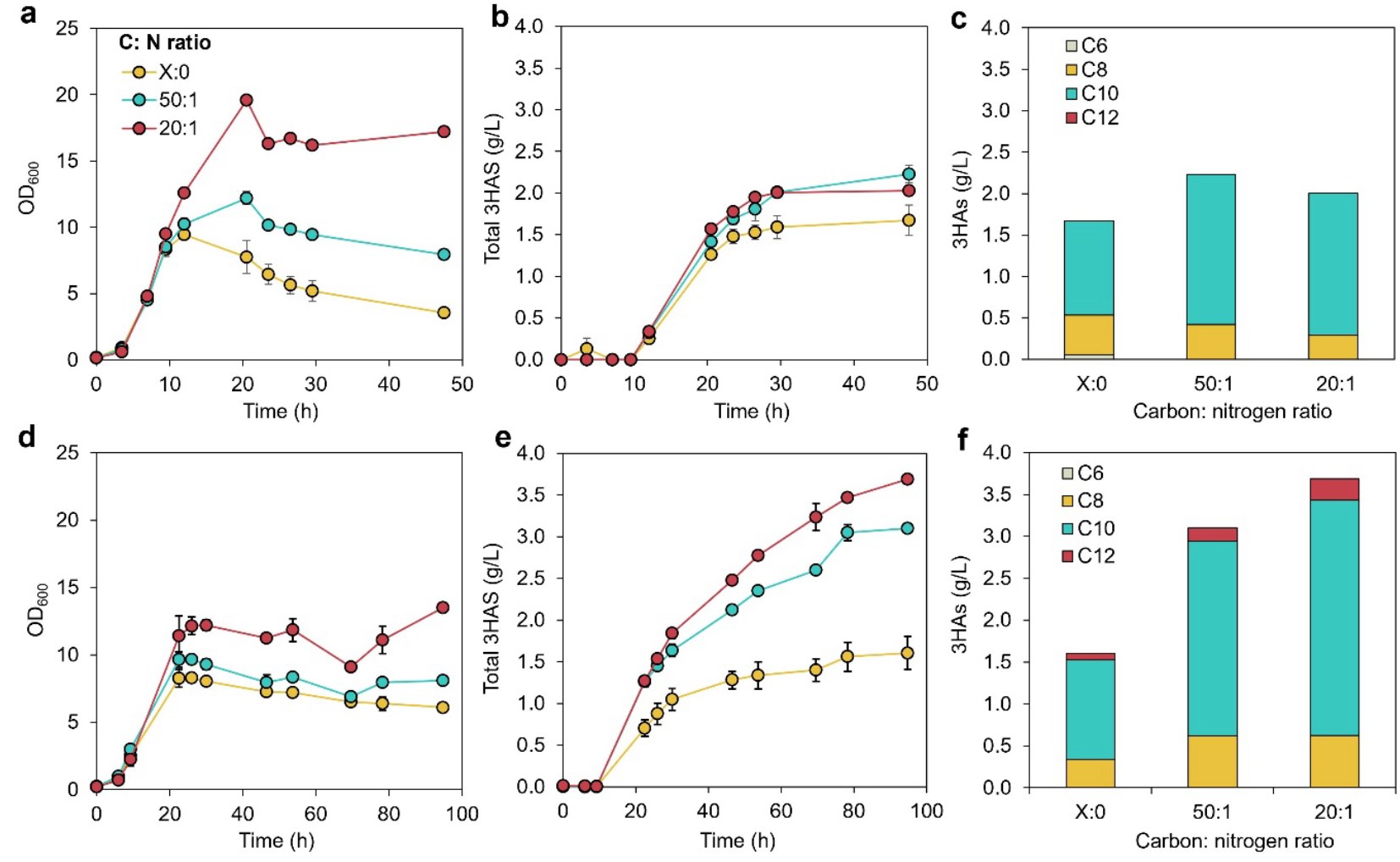
Cultivation profiles of *P. putida* mutant strains during 3-HA production from glucose in 0.5 L bioreactors. Carbon-to-nitrogen (C:N) molar ratios of 20:1, 50:1, and a case with no-nitrogen (X:0) were tested. Cultivation profiles from: **(a-c)** a DO-stat fed batch feeding strategy for *P. putida* Δ*alkK*Δ*fadD1D2*, and **(d-f)** a continuous feeding strategy for cultivations of *P. putida* Δ*alkK*Δ*fadD1D2*Δ*gcd*Δ*hex*. **(a, d)** Profiles for optical density (OD_600_), used as a proxy of bacterial growth, **(b, e)** production of total 3HA, and **(c, f)** composition of 3-HAs. Profiles for glucose remaining in the bioreactor, glucose added to the bioreactors, and 2-ketogluconate accumulation are shown in **Figures S6-S7**. Data points are the average of biological duplicates and error bars represent the absolute error between duplicates, except for the carbon-nitrogen (C:N) molar ratio of 20:1, which is a single data point.

To prevent loss of carbon from glucose in the form of 2-ketogluconate, and because 2-ketogluconate accumulation would be detrimental for 3-HA recovery, we knocked out *gcd* and *hexR* in the Δ*alkK*Δ*fadD1D2* strain to avoid its accumulation and accelerate growth.^31,32^ The resulting strain, Δ*alkK*Δ*fadD1D2*Δ*gcd*Δ*hexR*, was again evaluated for the production of 3-HA at different C:N ratios. Because the accumulation of 2-ketogluconate should not be an issue in this strain, we used a feeding mode to maintain glucose levels between 2-10 g/L. Similar to the DO-stat fed-batch cultivations, cell growth was the highest at a C:N ratio of 20 (**Figure 4d**). Production of 3-HA was improved in all cases except when nitrogen was not added to the feeding media (**Figure 4e**). At a C:N ratio of 20, the total 3-HA production was 3.7 g/L (from 55 g/L of glucose added, while a titer of 3.1 g/L was obtained at a C:N ratio of 50 (from 45 g/L of glucose) (**Figure 4e, Figure S7**). 3-HDA was again the most abundant 3-HA at approximately 75% in all conditions (mass basis) and in this condition, there was also measurable production of the C12 3-HA (5%) (**Figure 4f**).

## Discussion

In this work, we demonstrated that the traditional understanding of the PHA biosynthetic pathway in *P. putida* was incomplete, namely from *in vitro* biochemistry and *in vivo* gene deletion experiments, we conclude that PhaG hydrolyzes 3-hydroxyacyl-ACP to free 3-HA. Previous *in vitro* studies concluded PhaG performs 3-hydroxyacyl-ACP:CoA transferase activity; however, these studies only examined the reverse reaction and do not provide direct experimental evidence for 3-hydroxyacyl transfer from ACP to CoA.^9,23^ Here we assayed the forward reaction *in vitro*, showing PhaG catalyzes formation of 3-HDA from 3-hydroxydecanoyl-ACP in a stoichiometric ratio to precursors in the enzyme cascade. Furthermore, *in vivo* growth experiments presented here show that CoA ligases are required for PHA biosynthesis, aligning with *in vitro* assay results. In the absence of 3-HA-dependent CoA ligases, *P. putida* produces 3-HA instead of PHA, and polymerization requires the 3-HA to first be activated with CoA. Based on these findings, we propose a redefined PHA pathway where PhaG performs 3-HA-ACP hydrolysis and either FadD1 or AlkK performs CoA ligation (**Figure 1**) prior to polymerization by PhaC. This work not only improves our understanding of a critical pathway for bacterial survival in the environment but also can help guide metabolic engineering strategies for improved production of PHAs and related compounds such as 3-HA.

Previous efforts in designing biological production pathways of *mcl*-3-HA have primarily required PHA production followed by degradation into 3-HA either *in vivo* by PhaZ or chemically.^21,22^ Identifying that the conversion of 3-hydroxyacyl-ACP to 3-hydroxyacyl-CoA is performed in two steps, by separate enzymes, allowed us to design a direct biological synthesis route, where the PHA pathway is interrupted after 3-hydroxyacid-ACP hydrolysis by deletion of *fadD* and *alkK*. The Δ*alkK*Δ*fadD1D2* strain achieved a higher titer of 3-HA, (∼0.6 g/L) compared to PHA production titers (∼0.3 g/L) by the WT strain in comparable shake-flask cultures. This engineered pathway avoids unnecessary reactions that the cell needs to expend energy on, and 3-HA titers are not limited by cell volume, PHA production titers, and depolymerization rates. To further convert carbon from glucose into 3-HA production and prevent 2-ketogluconate accumulation, deletion of *gcd* and *hexR* enabled production of 3.7 g/L 3-HA in a bioreactor at a C:N ratio of 20, demonstrating that the strain is ready for further optimization and potential industrial applications. Notably, the extracellular nature of *mcl*-3-HAs provides a product recovery advantage over intracellular PHAs. For instance, the use of *in situ* product recovery methods, such as liquid–liquid extraction,^33^ represents a viable process engineering strategy for further bioprocess improvement.

This engineered pathway starts from fatty acid biosynthesis; thus, a variety of carbon sources can be used for 3-HA production, allowing flexibility to select an inexpensive feedstock. In this study we used glucose and *p*-coumarate as carbon sources, but this approach should be further expandable to include other low-cost feedstocks that can be consumed by wild type or engineered *P. putida* strains, such as waste glycerol, C5 sugars present in plant biomass,^34–36^ and plastics^37,38^. Additionally, because 3-HA are exported, this pathway may enable easier downstream processing to further optimize production relative to cell lysis and PHA recovery.

## Methods

### General strain cultivation

Plasmids and strains used in this study are listed in **Tables S2** and **S4**. Strains were grown in LB (Miller) broth (10 g/L tryptone, 5 g/L yeast extract, and 10 g/L NaCl) for routine culturing and preparation of competent cells. *P. putida* mutants are derived from parent strain AG5577,^39^ which is derived from KT2440 and is metabolically wild type, with “landing pads” inserted into neutral loci to enable site-specific DNA integration into the chromosome via Serine recombinase-Assisted Genome Engineering (SAGE).^40^ Here, we treat AG5577 as a functionally wild type strain. *P. putida* cultures were grown in a modified version of M9 that contained: 6 g/L Na_2_HPO_4_, 3 g/L KH_2_PO_4_, 0.5 g/L NaCl, 100 µM CaCl_2_, 18 µM FeSO_4_, 2 mM MgSO_4_, and 1X MME trace minerals solution, with variable nitrogen and carbon concentrations as described in the below sections for PHA and 3-HA production. Prior to inoculation, media were sterilized either by autoclaving (LB) or filtration with a 0.22 µm filter (modified M9). Antibiotics for plasmid selection and maintenance were used at the following final concentrations: 50 µg/mL kanamycin, 50 µg/mL apramycin, and 100 µg/mL ampicillin, with the addition of agar (15 g/L) for growth on petri plates. Unless otherwise indicated, cultures were incubated at 37°C for *E. coli* or 30°C for *P. putida*. For maintenance of temperature sensitive plasmids, cultures were incubated at room temperature (approximately 22°C). Unless otherwise indicated, liquid cultures were agitated by shaking at 225 rpm with a 2.5 cm orbital in an Innova S44i (Eppendorf) incubator.

### Plasmid construction

Plasmids and oligonucleotides used in this study are listed in **Tables S2** and **S3**, respectively. Plasmids were constructed using enzymes from New England Biolabs (NEB) following the manufacturer’s recommended protocol. Phusion® or Q5® High-Fidelity Polymerases (NEB) were used for all PCR amplifications followed by DpnI (NEB) digestion to remove plasmid templates from PCR amplified products. DNA fragments were assembled with NEBuilder® HiFi DNA Assembly Master Mix (NEB) to produce the desired plasmid constructs. Plasmids were propagated in NEB 5-α F’*I*^*q*^ *E. coli* grown in LB (Miller) supplemented with the appropriate antibiotics for plasmid selection and maintenance (50 µg/mL kanamycin, 50 µg/mL apramycin, or 100 µg/mL ampicillin). Plasmids were extracted from overnight cultures using the geneJET Plasmid Miniprep Kit (Thermo Fisher Scientific).

### Protein production

Proteins used for the *in vitro* assay were heterologously produced in *E. coli* BL21(DE3) transformed with appropriate plasmids as listed in **Table S2**. Strains producing *Ec*ACP, *Ec*FabD, *Pp*FabH2, and *Pp*FabG were grown in LB (Miller) supplemented with 100 µg/mL ampicillin. An overnight starter culture was used to inoculate the main culture at 1% (v/v) and grown at 37°C at 200 rpm. When the OD_600_ reached ∼0.7, the culture was induced with 0.5 mM IPTG; and incubated for another ∼16 hours at 18°C, 200 rpm. To facilitate the post-translational modification of *Ec*ACP, this protein was co-produced with *Ec*AcpS and was supplemented with 0.1 mM of pantothenic acid at the time of induction.^41^ The strain producing *Pp*PhaG was co-transformed with pGro7 (Takara Bio Inc.) to produce the chaperones GroEL and GroES and was cultivated in LB (Miller) supplemented with 100 µg/mL ampicillin and 30 µg/mL chloramphenicol. This culture was supplemented with 0.1 mg/mL L-arabinose at the point of inoculation to induce chaperone production, but otherwise was grown and induced similarly as others. The resulting biomass was harvested by centrifugation and frozen at -80°C until purification.

*Ec*ACP, *Ec*FabD, *Pp*FabH2, and *Pp*FabG were purified following a combination of immobilized-metal affinity chromatography (IMAC) and anion exchange chromatography. The anion exchange chromatography was performed using an AKTA Pure fast protein liquid chromatography (FPLC) system (Cytiva). Protein purification was performed with Buffer A (25 mM HEPES, 100 mM NaCl at pH 7.5). The thawed biomass, supplemented with trace amounts of DNaseI, was lysed either by sonication or using a French press. The lysate was cleared by centrifugation and the supernatant filtered through a 0.45 μm filter before being applied to a gravity-fed column packed with Ni-sepharose resin or a HisTrap column (Cytiva). The column was washed with up to 20 mM imidazole and eluted with 200 mM imidazole. The eluate was concentrated using a centrifugal spin column with the appropriate size cutoff and further purified on a column packed with Source-15Q resin. The protein was eluted following an increasing gradient of NaCl in Buffer A. Fractions containing the protein of interest were pooled and concentrated using a centrifugal spin filter. Due to its inherent instability, *Pp*PhaG was only purified using IMAC with the purification buffer supplemented with 1 mM tris(2-carboxyethyl)phosphine (TCEP) and 10% (v/v) glycerol. All concentrated protein was frozen as beads over a bath of liquid N_2_ and kept at -80°C until further use.

### *In vitro* activity assays

The *in vitro* assay for determining the activity of *Pp*PhaG was performed in a one-pot reaction and the resulting reaction product was detected using LC-MS. The assay was performed at room temperature in 50 mM HEPES, pH 7.5 supplemented with 10 mM MgCl_2_ and 1 mM TCEP. The reaction was initiated by mixing 200 μM of *Ec*ACP, 5 μM *Ec*FabD, and 500 μM malonyl-CoA (Sigma-Aldrich). After 5 minutes, the reaction was supplemented with 5 μM *Pp*FabH2 and 500 μM octanoyl-CoA (Cayman Chemical). Following another 5-minute incubation, the mixture was supplemented with 5 μM *Pp*FabG and 500 μM NADPH and incubated for 5 minutes. Lastly, the reaction was supplemented with ∼1 mg/mL partially purified *Pp*PhaG and incubated for 15 minutes. Equivalent reactions devoid of *Pp*PhaG were also analyzed for comparison. The mixture was quenched with 5% (v/v) glacial acetic acid and the precipitated protein was removed using a combination of centrifugation and 0.2 μm filtration prior to LC-MS analysis. Three independent reactions with and without *Pp*PhaG were analyzed in this study.

### LC-MS method analysis of *in vitro* assay

Quantitation of samples was performed on an Agilent 1100 LC system (Agilent Technologies) equipped with an Agilent G6120A single quadrupole mass spectrometer. Each sample and standard were injected at a volume of 10 μL on a Phenomenex Luna C18(2)-HST 100Å column (25 μm, 2.0 × 100 mm). The column temperature was maintained at 45°C and the buffers used to separate the analytes of interest were: 0.16% formic acid in water (A) and acetonitrile (B). The following gradient method was used to separate the analytes of interest: 0-1 min, 0% B; 1-7.67 min, 0% to 50% B; 7.67-9.33 min, 50% to 70% B; 9.33-10.67 min, 70% B; 10.67-10.68 min, 70% to 100% B; 10.68-11.68 min, 100% B; 11.68-11.69 min, 100% to 0% B; 11.69-14 min, 0% B. The flow rate was held constant at 0.50 mL min^-1^, resulting in a run time of 14 minutes. The mass spectrometer was scanned in positive electrospray ionization mode from 50-100 *m/z* with a gas temperature of 350°C, drying gas flow at 12.0 L/min, nebulizer pressure of 35 psig, and a VCap of 3000v. The total ion chromatogram (TIC) and retention time were used to identify and quantitate the analyte of interest. A 3-hydroxydecanoic acid standard (Sigma-Aldrich) was used to construct a calibration curve between the ranges of 25 – 100 μg/mL. A minimum of 4 calibration levels were used for 3-hydroxydecanoic acid with an r^2^ coefficient of 0.995 or better. A check calibration standard (CCS) was analyzed every 10 samples to ensure the integrity of the initial calibration.

### Transcriptomics analysis

*P. putida* strains were grown in modified M9 medium with 25 mM acetate or 5 mM 3-HDA as the sole carbon source. Cultures were inoculated at a starting OD_600_ of 0.1 and maintained at 30°C, 230 rpm until reaching OD_600_ 0.3. RNAprotect Bacteria Reagent (Qiagen) was directly added to the culture and cells were harvested by centrifugation and sent to CD Genomics for RNA extraction and sequencing. Reads were mapped to the *P. putida* KT2440 genome (Accession number NC_002947) using Kallisto^42^. Transcript abundance and differential gene expression were calculated with Voom and Limma using Degust^43^. Relative abundance of RNA for each CoA ligase in the 3-hydroxydecanoate growth condition compared to acetate was ranked. A greater than zero log_2_ fold change indicates an increase in transcript abundance and a less than zero indicates a decrease. RNA-seq datasets have been deposited in the NCBI Gene Expression Omnibus (GEO) under accession number: GSE12188.

### *P. putida* mutant strain construction

Gene deletions were carried out with pK18mobsacB-derived plasmids using a previously described kanamycin selection and sucrose counter selection protocol.^44,45^ *FadD1* complementation was achieved following the previously described Bxb1 serine integrase mediated genome integration and ϕC31 plasmid backbone excision protocol.^39,40^ To prepare *P. putida* electrocompetent cells, glycerol stocks were used to inoculate 50 mL LB (Miller) in a 250 mL shake flask, and cultures were incubated for 16-18 hours at 30°C, shaking at 225 rpm. Cells were collected by centrifugation at 5000 × *g* for 10 minutes at room temperature (approximately 22°C), and washed three times with 25 mL of 10% glycerol followed by a final resuspension in 1 mL of 10% glycerol. Aliquots were stored at -80°C until further use. Electroporation was performed with 1 mm gap cuvettes containing 50 μL electrocompetent *P. putida* cells and either 300-800 ng of gene deletion plasmids or 20 ng each of Bxb1 integrase and cargo plasmids. The mixture was electroporated using a Gene Pulser Xcell (Bio-Rad) set to 1600 V, 25 µF, 200 ohm and recovered with 950 μL SOC outgrowth medium (NEB) for 1 hour at 30°C, 225 rpm. Gene integrations and deletions were confirmed by PCR.

### Growth experiments of *P. putida* on 3-hydroxyacids as the sole carbon source

To prepare seed cultures, glycerol stocks of *P. putida* strains were used to inoculate 5 mL LB broth (Miller) and incubated overnight at 30°C, 225 rpm. A 1% (v/v) inoculum from the overnight culture was used to inoculate a second seed culture of 5 mL modified M9 medium containing 2 mM (NH_4_)_2_SO_4_ and 5 mM glucose, which was then incubated overnight at 30°C, 225 rpm. Next, 1% inocula from the second seed cultures were transferred to a 48-well clear, flat bottom, cell culture plate (Greiner, CELLSTAR®) containing 0.5 mL modified M9 with 2 mM (NH_4_)_2_SO_4_ and either 5 mM 3-hydroxydecanoic acid or 3-hydroxyoctanoic acid. Plates were incubated in a BioTek Epoch2 microplate reader at 30°C, continuous fast shaking at 548 cpm, in double orbital mode (2 mm), and absorbance at 600 nm was recorded at 10-minute intervals. All cultures were grown in triplicate.

### Sample preparation cryogenic electron microscopy

Bacterial samples were prepared for cryo-electron microscopy (cryo-EM) analysis by applying 3 µL of prepared bacterial suspension to glow discharged Quantifoil® Copper 2/1 grids (300 mesh). The glow discharging was performed using a GloQube™ (EMS) plasma cleaner at 5 mA for 15 seconds. Grids were plunge frozen in nitrogen-cooled ethane using a Vitrobot™ Mark IV (ThermoFisher Scientific) equilibrated to 100% humidity and 4°C with a blotting time of 3 seconds and a blot force of -4.

### Cryo-EM data collection and processing

Cryo-EM data were collected using a ThermoFisher Scientific Krios™ G4 transmission electron microscope operated under cryogenic conditions. Imaging conditions were as follows: a nominal magnification of 14,000X in TEM nanoprobe mode with a parallel beam, spot size 8, an illuminated area of 3.57µm, a constant defocus of -5µm, and a 70µm C2 aperture. Data were collected with a Falcon™ 3EC direct electron detector (ThermoFisher Scientific) in linear mode. The total electron dose was limited to 60 e/Å^2^ to minimize radiation damage. Collected movies were corrected for sample and stage drift using the Patch Motion Correction option in CryoSPARC™.

### Cultivation and sample preparation for PHA production from shake flask experiments

Culture flasks with 5 mL LB (Miller) medium were inoculated from glycerol stocks, and incubated for 16-18 hours at 30°C, 250 rpm. In triplicate, 5 µL of the overnight LB cultures were used to inoculate 5 mL modified M9 medium with 2 mM (NH_4_)_2_ SO_4_ and 25 mM *p*-coumarate as the sole carbon source. Cultures were incubated for 16-18 hours at 30°C, 250 rpm.

Subsequently, 500 µL of each overnight cultures were used to inoculate 50 mL of the same modified M9 medium in 250 mL Erlenmeyer flasks and incubated at 30°C, shaking at 250 rpm for 72 hours. Cells were harvested by centrifugation, pellets were washed twice with deionized water and lyophilized.

### *mcl*-PHA composition analysis via derivatization and gas chromatography-mass spectrometry (GC-MS)

Lyophilized samples were then analyzed by GC-MS post depolymerization and derivatization to hydroxyacyl methyl esters (HAMEs), as previously reported,^3^ noting a change of the GC column and supplier to a Restek Stabilwax-DA column (30 m × 0.25-mm id, 0.25-μm film).

### Cultivation and sample preparation for HPLC analysis of 3-HA production from shake flask experiments

Cultures were prepared as described in the PHA production section. Strains were grown in modified M9 with various carbon sources and 2 mM (NH_4_)_2_SO_4_ to induce 3-HA production. Cultures were incubated at 30°C shaking at 250 rpm for up to 96 hours, and 500 µL samples were periodically collected for HPLC analysis. Samples were centrifuged, and the supernatant was acidified with H_2_SO_4_ and filtered with a 0.22 μm spin filter. Samples were analyzed using a 1260 Infinity II HPLC (Agilent), equipped with a Fast-Acid column (Bio-Rad) heated to 60°C, and monitored with a refractive index detector (RID), maintained at 35°C. 5 µL aliquots of each sample were resolved using an isocratic elution of 5 mM H_2_SO_4_ in H_2_O, and the RID was monitored for 3-HA. Flow rate was maintained at 0.6 mL/minute. Standard curves were generated using Agilent OpenLab CDS for 3-hydroxyhexanoate, 3-hydroxyoctanoate, and 3-hydroxydecanoate (Sigma Aldrich) and carbon sources used in the culture fermentation.

### Analysis of 3-HA production from bioreactor experiments by UHPLC-MS/MS operating in multiple reaction monitoring (MRM) mode

Even chain 3-hydroxy acids (3-HAs) ranging from C6 to C12 were quantified using ultra-high performance liquid chromatography coupled with tandem mass spectrometry (UHPLC-ESI-MS/MS) operating in multiple reaction monitoring (MRM) mode. The method enables sensitive and selective quantitation of 3-hydroxyhexanoic, 3-hydroxyoctanoic, 3-hydroxydecanoic, and 3-hydroxydodecanoic acids in biological samples.

#### Standard Preparation

Even chain 3-hydroxy acid standards (C6–C12) were purchased from Matreya (now a part of Cayman Chemicals). Individual stock solutions were prepared at 1 mg/mL in methanol and combined to form a working standard solution of 100 µg/mL. Calibration standards were prepared by diluting the working solution to produce concentrations ranging from 0.5 µg/mL to 100 µg/mL with a minimum of 6 points, and a calibration verification standard at 20 µg/mL.

#### Chromatography

Chromatographic separation was performed on an Agilent 1290 Infinity UHPLC system using a ZORBAX RRHD Eclipse Plus C18 column (2.1 × 50 mm, 1.8 µm), and the column temperature was maintained at 35 °C. The mobile phase was delivered at a constant flow rate of 0.5 mL/min, using the following solvents: (A) 0.1% formic acid with 5 mM ammonium formate in water, and (b) 0.1% formic acid with 5 mM ammonium formate in methanol. The gradient began with 99% solvent A and 1% solvent B for the first minute, followed by a linear increase in solvent B to 99% over the next six minutes. At 7.0 minutes, the gradient was returned to the initial conditions of 100% A, and the system was re-equilibrated under these conditions from 7.01 to 9.0 minutes before the next injection. Samples and standards were injected at a volume was 0.5 µL.

#### Mass Spectrometry

Analytes were detected using an Agilent 6470A triple quadrupole mass spectrometer equipped with an Agilent Jet Stream electrospray ionization (AJS-ESI) source, which was operated in negative ion mode. The ion source was configured with a gas temperature of 300 °C and a gas flow rate of 10 L/min, while the nebulizer pressure was set to 25 psi. The sheath gas was maintained at a temperature of 375 °C with a flow rate of 12 L/min. The capillary and nozzle voltages were set to 4500 V and 500 V, respectively. Prior to analysis, multiple reaction monitoring (MRM) transitions for all compounds were optimized. The quantifying and qualifying transitions, along with their corresponding collision energies and fragmentor voltages, are summarized in **Table S5**. Quantitation was performed using external standard calibration curves generated from peak areas of the quantifying transitions. Calibration curves were fitted using a quadratic model and exhibited correlation coefficients (r^2^) ≥ 0.995.

Data acquisition and processing were conducted using Agilent MassHunter Quantitative Analysis (QQQ) software, version 10.2.

### Seed train for bioreactor cultivations

Engineered strains were maintained at -80°C in 20% glycerol (v/v). The surface of the glycerol stocks was scraped with a sterile loop and used to inoculate 50 mL of LB (Miller) medium in 250 mL flasks. These seed cultivations were incubated at 30°C and 180 rpm for 16 h. After this time, cells were centrifuged for 5 min at 5000 × *g* and resuspended in 5 mL of a modified version of M9 medium that contained: 13.56 g/L Na_2_HPO_4_, 6 g/L KH_2_PO_4_, 1 g/L NaCl, 2 g/L (NH_4_)_2_SO_4_, 2 mM MgSO_4_, 100 µM CaCl_2_ and 36 µM FeSO_4_.

### General methodology for cultivations performed in 0.5-L bioreactors

All bioreactor cultivations were performed in 0.5-L Biostat-Q Plus system bioreactors (Sartorius Stedim Biotech). Unless otherwise stated, the batch medium for each experiment (250 mL) consisted of modified M9 medium supplemented with 10 g/L glucose. Three drops of sterile Antifoam 204 (Sigma Aldrich, United States) were added before inoculation. The bioreactors were sparged with air at 300 mL/min to reach an initial dissolved oxygen (DO) saturation of 100% with a starting agitation of 350 rpm. Bioreactors were inoculated at an initial optical density at 600 nm (OD_600_) of 0.2 (measured with Nanodrop™, Thermo Scientific). Cultivations were maintained at 30°C and pH 7, adjusting with 4 N NaOH as necessary. Using sterile technique, samples were collected to measure bacterial growth (OD_600_), concentration of sugars, and production of metabolites. Cultivations were generally ended due to excessive foaming, at 48 h for DO-stat fed batch, and at 96 h for continuous feeding fed-batch.

### DO-stat fed-batch cultivations in bioreactors

We selected a DO-stat fed-batch strategy to evaluate the effect of different carbon to nitrogen ratios on the production of 3-HAs. This strategy was only used for cultivations with the Δ*alkK*Δ*fadD1D2* strain, AG7303. The batch phase aimed to generate microbial biomass, and the fed-batch phase aimed to produce 3-HAs. The feeding medium consisted of modified M9 with 500 g/L glucose and (NH_4_)_2_SO_4_ concentrations of 55.1 g/L, 22 g/L, or 0 g/L, which correspond to carbon atom to nitrogen atom ratios of 20, 50, and infinity on a molar basis, respectively. The solutions were adjusted to pH 7 with 4 N NaOH. Once the batch growth phase was complete, the feeding solution was pumped for 30 seconds to feed ∼1 mM glucose per pulse every time the DO reached 75%. The agitation was manually adjusted (**Figure S5**). Cultivations were concluded when DO did not change after the addition of feed media. Antifoam was added as needed during foaming events. All bioreactor runs were performed in duplicate.

### Fed-batch cultivations in bioreactors with continuous feeding

Fed-batch cultivations with continuous feeding were performed with the Δ*alkK*Δ*fadD1D2*Δ*gcd*Δ*hexR* strain, AG10129. After inoculation, DO was allowed to drop to 30%, and it was automatically controlled at that level by automatic changes in the agitation speed. The feeding medium was the same as the one described above, with the same glucose-to-nitrogen ratios. After the batch growth phase was complete, the feeding rate was initiated at 0.83 mL/h (1.66 g/L/h glucose), and the rate was adjusted manually to control glucose at concentrations below 10 g/L. The cultivation ended at 96 h. Antifoam 204 was added as needed during foaming events.

## Data availability

RNA-seq datasets have been deposited in the NCBI Gene Expression Omnibus (GEO) under accession number: GSE12188. Source data for growth curves, 3-HA and PHA production are provided as a Source Data file. Plasmid maps are provided in the Supplemental Information file.

## Acknowledgements

Kevin J. McNaught is thanked for the construction of pKM175 and pKM181. Joshua Elmore is thanked for the construction of pJE1369. Sean P. Woodworth is thanked for providing PHA analysis via GC-MS.

Electron microscopy was conducted at the Center for Nanophase Materials Sciences, which is a US Department of Energy Office of Science User Facility at Oak Ridge National Laboratory.

This work was authored in part by Oak Ridge National Laboratory, which is managed by UT-Battelle, LLC, for the U.S. Department of Energy (DOE) under contract DE-AC05-00OR22725. This work was also authored by the National Renewable Energy Laboratory, operated by Alliance for Sustainable Energy, LLC, for the U.S. Department of Energy (DOE) under Contract No. DE-AC36-08GO28308. Funding was provided by the U.S. Department of Energy, Office of Energy Efficiency and Renewable Energy, Bioenergy Technologies Office (BETO) and Advanced Materials and Manufacturing Technologies Office (AMMTO) as part of the BOTTLE Consortium. Funding was also provided in part by BETO for the Agile BioFoundry. The views expressed in the article do not necessarily represent the views of the DOE or the U.S. Government. The U.S. Government retains and the publisher, by accepting the article for publication, acknowledges that the U.S. Government retains a nonexclusive, paid-up, irrevocable, worldwide license to publish or reproduce the published form of this work, or allow others to do so, for U.S. Government purposes.

