## Supplemental Figures and Tables for "The redefined role of PhaG and CoA ligases in medium-chain-length 3-hydroxy acid and polyhydroxyalkanoate production in *Pseudomonas putida*"

### Authors & Affiliations

A. R. Ola Pasternak,[1,2] Walter T. Woodside,[1,2] Eugene Kuatsjah,[2,3] Sekgetho C. Mokwatlo,[2,3] Jay D. Huenemann,[1,2] William E. Michener,[2,3] Stefan J. Haugen,[2,3] Darren J. Parker,[1,2] Alexis N. Williams,[4] Kelsey J. Ramirez,[2,3] Davinia Salvachúa,[2,3] Gregg T. Beckham,[2,3*] and Adam M. Guss[1,2,*]

[1] Biosciences Division, Oak Ridge National Laboratory, Oak Ridge, TN, 37831 USA.

[2] BOTTLE Consortium, National Renewable Energy Lab, USA.

[3] Renewable Resources and Enabling Sciences Center, National Renewable Energy Laboratory, Golden, CO 80401

[4] Center for Nanophase Materials Sciences, Oak Ridge National Laboratory, Oak Ridge, TN 37830, USA

Notice: This manuscript has been authored by UT-Battelle, LLC, under Contract No. DE-AC0500OR22725 with the U.S. Department of Energy. The United States Government retains and the publisher, by accepting the article for publication, acknowledges that the United States Government retains a non-exclusive, paid-up, irrevocable, world-wide license to publish or reproduce the published form of this manuscript, or allow others to do so, for the United States Government purposes. The Department of Energy will provide public access to these results of federally sponsored research in accordance with the DOE Public Access Plan (http://energy.gov/downloads/doe-public-access-plan).

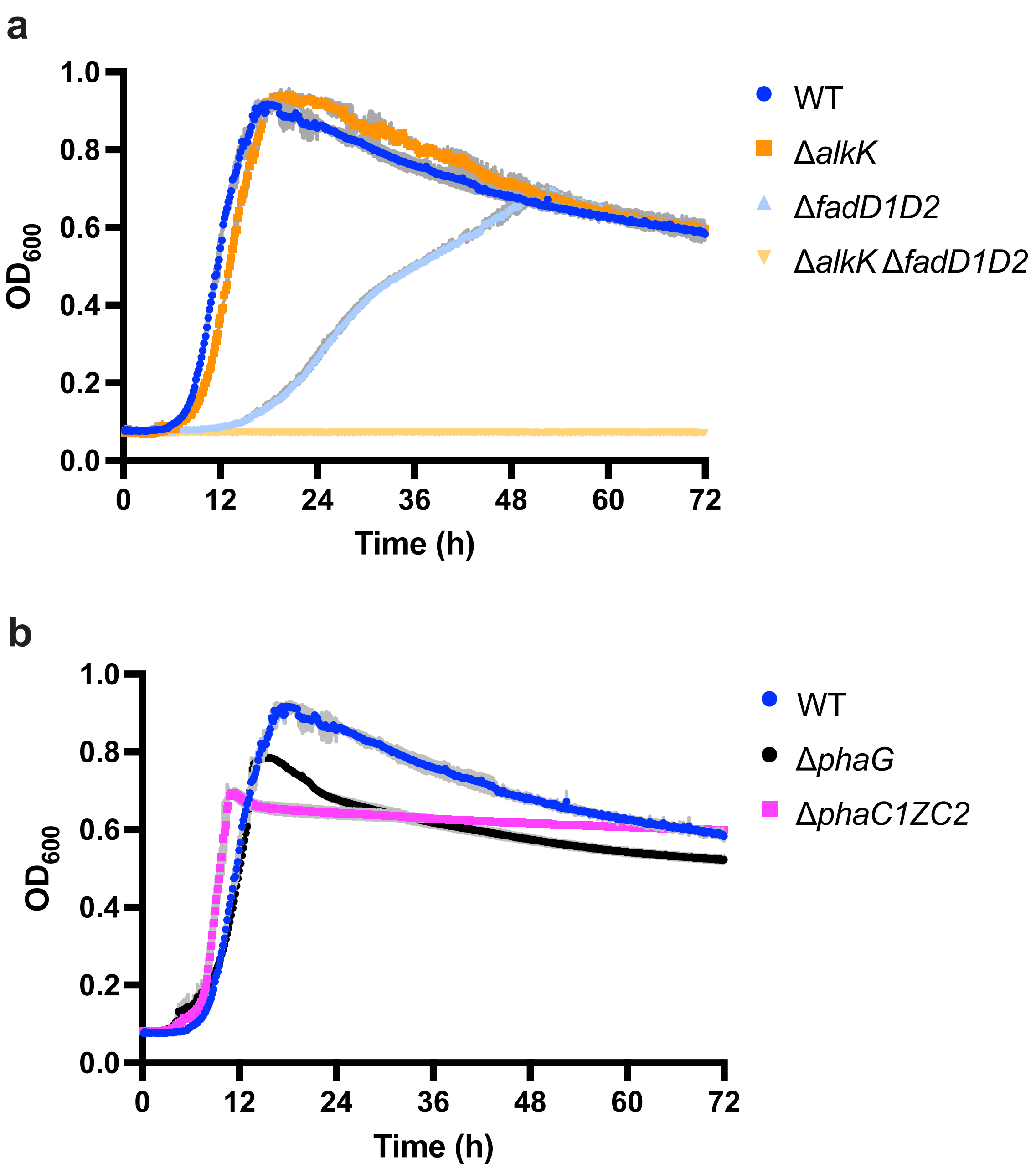


**Figure S1. *P. putida* Δ*alkK* and Δ*phaG* strains grow similarly to the parent strain on 3-HDA.** Growth curves of *P. putida* mutants on modified M9 medium with 3-hydroxydecanoic acid as the sole carbon source. a) *P. putida* parent strain AG5577 (dark blue), *P. putida* Δ*alkK* (orange), *P. putida* Δ*fadD1D2* (light blue), *P. putida* Δ*alkK* Δ*fadD1D2* (yellow). b) *P. putida* parent strain AG5577 (dark blue), *P. putida* Δ*phaG* (black), *P. putida* Δ*phaC1ZC2* (magenta). Data are represented as the average of three biological replicates with error bars showing standard deviation. Source data are provided as a Source Data file.

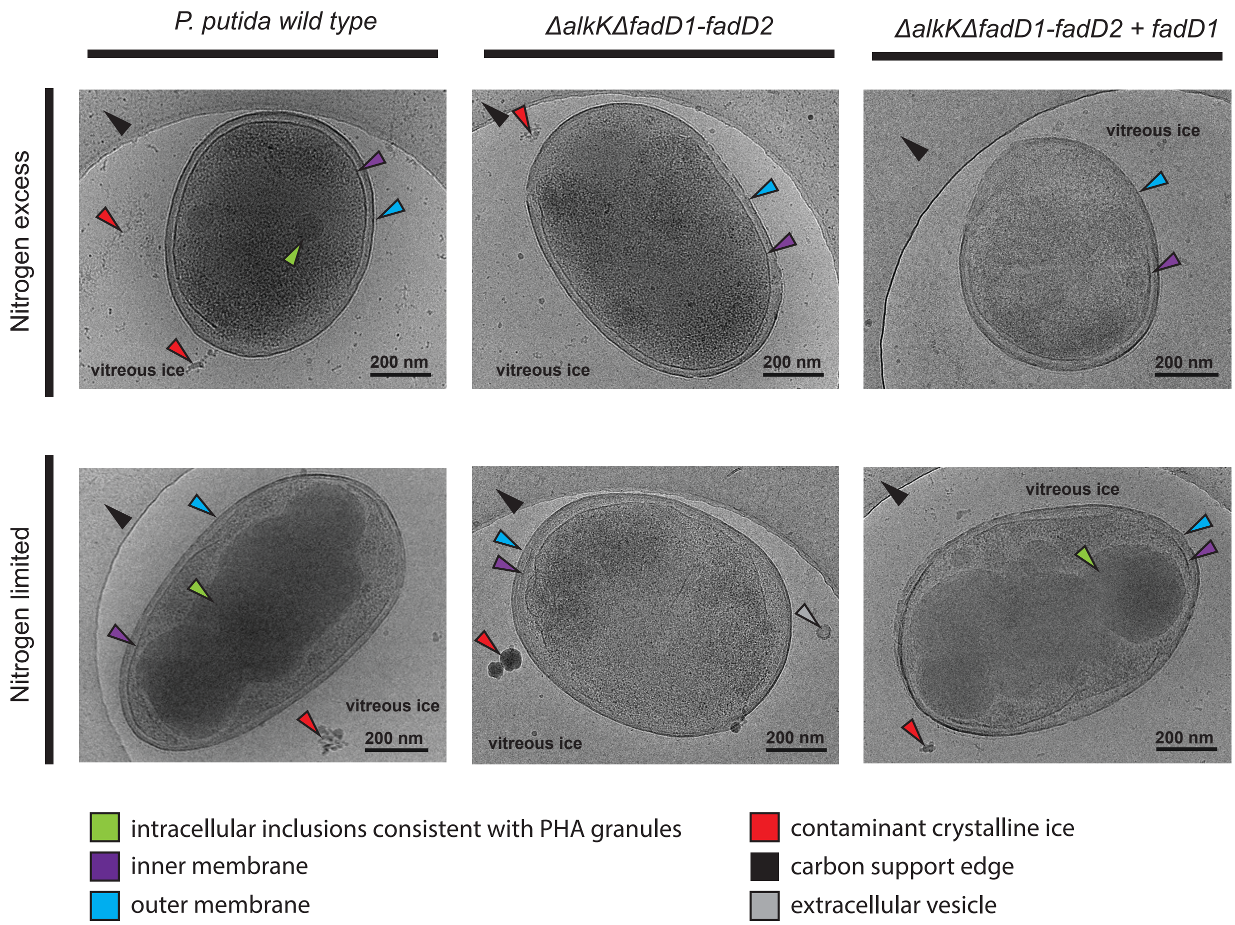


**Figure S2: Cryogenic electron microscopy images of *P. putida* strains grown in nitrogen-excess and nitrogen-limited conditions.** Features of interest highlighted with the following arrows: intracellular inclusions consistent with PHA granules (green), outer membrane (cyan), inner membrane (purple), vitreous ice (black text), contaminant crystalline ice (red), extracellular vesicle (grey), carbon support edge (black), scale bars: 200 nm.

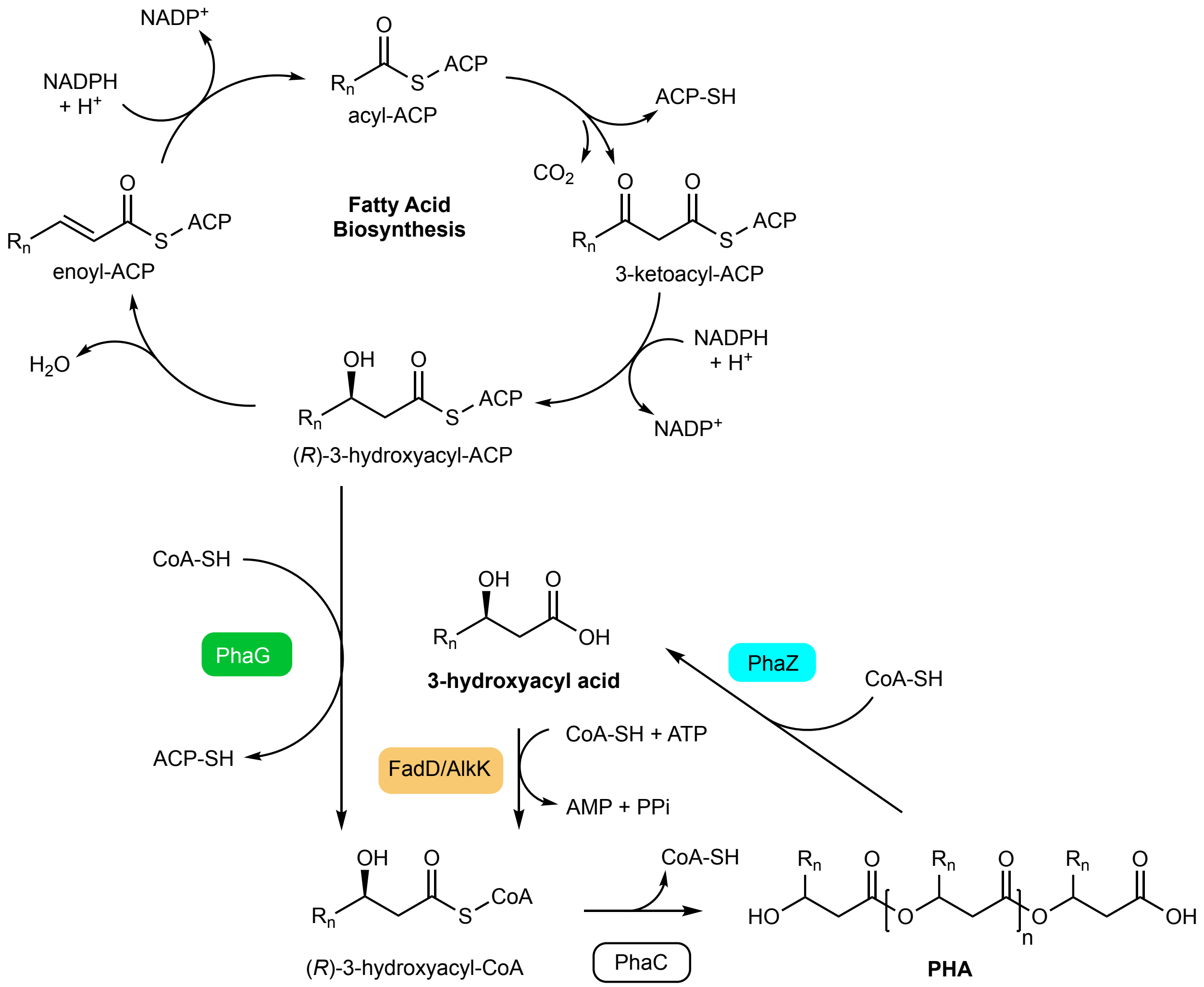


**Figure S3: Alternative hypothetical PHA biosynthetic pathway.** In this potential pathway, PhaG is a CoA transferase, and free 3-hydroxyacids are produced through hydrolysis of PHA by PhaZ. In this scenario, deletion of *phaZ* would prevent the accumulation of free 3-hydroxyacyl acids.

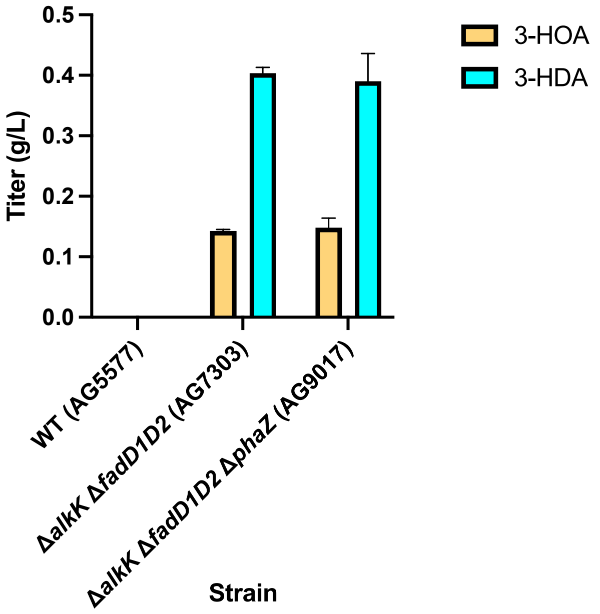


**Figure S4: Deletion of *phaZ* does not impact 3-HA accumulation.** Product titers of 3-hydroxyoctanoate (yellow) and 3-hydroxydecanoate (cyan) in *P. putida* mutant strains, n=3 when grown in shake flasks with 25 mM *p*-coumarate under N-limited conditions. Source data are provided as a Source Data file.

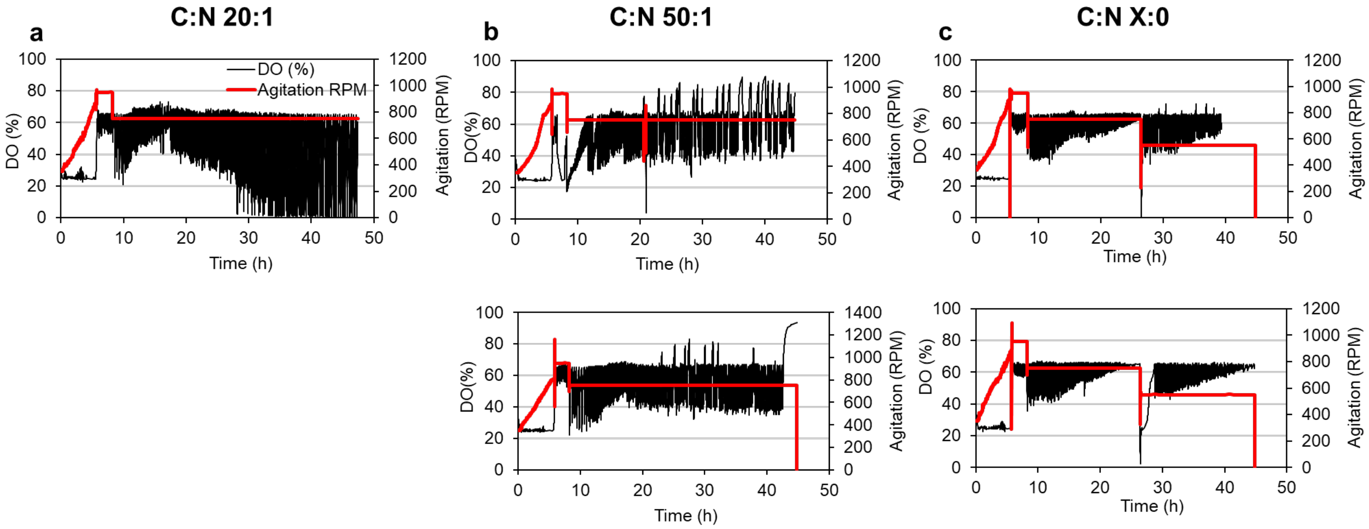


**Figure S5. Agitation and dissolved oxygen (DO) profiles of *P. putida* Δ*alkK*Δ*fadD1D2* (AG7303) during the production of 3-HA in 0.5-L bioreactors at different carbon-to-nitrogen ratios. (a)** Profile for C:N ratio of 20 (singlet), **(b)** profiles for C:N ratio of 50 (two replicates), and **(c)** profiles for C:N ratio of X:0 (two replicates).

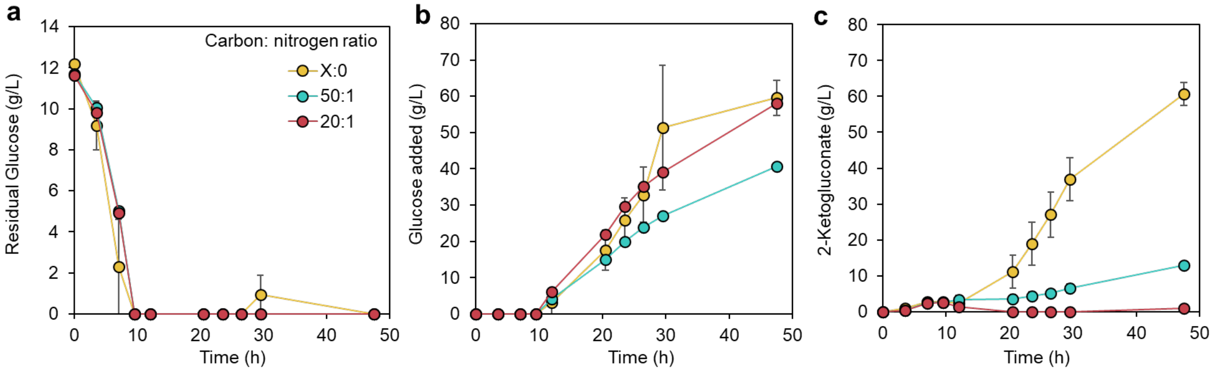


**Figure S6: Cultivation profiles of *P. putida* Δ*alkK*Δ*fadD1D2* during the production of 3-HA in 0.5 L bioreactors at different carbon-to-nitrogen ratios. (a)** Profiles for residual glucose in the bioreactors, **(b)** glucose added to the bioreactors, and **(c)** 2-ketogluconate accumulation in the bioreactor. Data points are the average of biological duplicates and error bars represent the absolute error between duplicates, except for carbon carbon-nitrogen ratio of 20:1, which is a singlet.

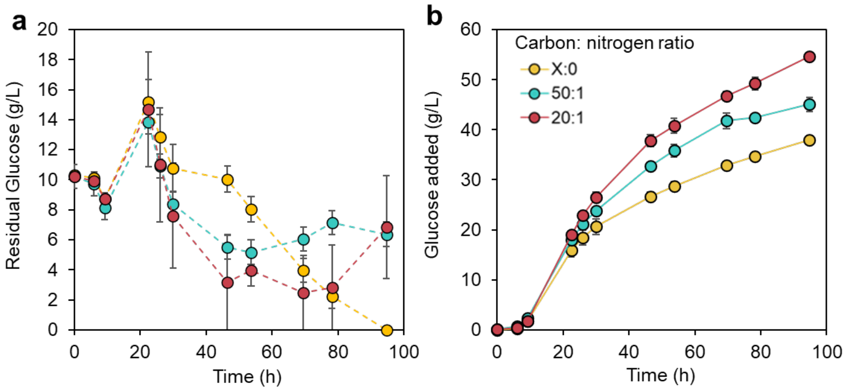


**Figure S7: Cultivation profiles of *P. putida* Δ*alkK*Δ*fadD1D2*Δ*gcd*Δ*hexR* during the production of 3-HA in 0.5 L bioreactors at different carbon-to-nitrogen ratios. (a)** Profiles for residual glucose in the bioreactors and **(b)** glucose added to the bioreactors. Data points are the average of biological duplicates and error bars represent the absolute error between duplicates.

### Supplementary tables

**Table S1. Transcriptomics analysis of wild type *P. putida* grown on 3-hydroxydecanoate relative to acetate.**

| *P. putida* fatty acid CoA ligase | $Log_2$ fold change | P-value |
|---|---|---|
| PP_4549 (*fadD1*) | 3.3508 | 9.02E-13 |
| PP_0763 (*alkK*) | 0.7888 | 0.0001235 |
| PP_3015 | 0.6865 | 0.215909 |
| PP_4063 | 0.3783 | 0.0809324 |
| PP_3279 (*paaK*) | 0.1307 | 0.562815 |
| PP_2038 | -0.1250 | 0.81353529 |
| PP_2709 | -0.2316 | 0.3481405 |
| PP_4550 (*fadD2*) | -0.3018 | 0.04560269 |
| PP_2213 | -0.3370 | 0.1541423 |
| PP_3553 | -0.5717 | 0.0071052 |
| PP_2766 | -1.8626 | 9.32E-06 |
| PP_2351 (*prpE*) | -2.1591 | 2.52E-09 |
| PP_4487 | -3.8293 | 6.99E-15 |

**Table S2. Plasmids used in this study.**

| Plasmid | Utility | Construction details |
|---|---|---|
| pKM175 | Heterologous production of *Ec*AcpP (b1094) in *E. coli* | pETDuet-1 T7 expression vector containing b1094 amplified from *E. coli* MG1655 gDNA (oKM0278 & oKM0279) in MCS-1 between insertion points: BamHI and EcoRI. |
| pKM181 | Heterologous production of holo-*Ec*AcpP (b1094), in conjunction with *Ec*AcpS (b2563), in *E. coli* | pETDuet-1 T7 expression vector derived from pKM175 to contain b2563 amplified from *E. coli* MG1655 gDNA (oKM0280 & oKM0281) in MCS-2 between insertion points: NdeI and XhoI. |
| pEUK070 | Heterologous production of *Ec*FabD (P0AAI9) in *E. coli* | pET15b(+) T7 expression vector containing P0AAI9 amplified from MG1655 gDNA (oEUK116 & oEUK117) using Gibson assembly between insertion points: NdeI and BamHI. |
| pEUK079 | Heterologous production of *Pp*FabH2 (PP_4545) in *E. coli* | pET15b(+) T7 expression vector containing PP_4545 amplified from KT2440 gDNA (oEUK128 & oEUK129) using Gibson assembly between insertion points: NdeI and BamHI. |
| pEUK086 | Heterologous production of *Pp*PhaG (PP_1408) in *E. coli* | pET21b(+) T7 expression vector containing PP_1408 amplified from KT2440 gDNA (oEUK174 & oEUK175) using Gibson assembly between insertion points: NdeI and BamHI. |
| pEUK087 | Heterologous production of *Pp*FabG (PP_1914) in *E. coli* | pET21b(+) T7 expression vector containing PP_1914 amplified from KT2440 gDNA (oEUK172 & oEUK173) using Gibson assembly between insertion points: NdeI and BamHI. |
| pGro7 | Plasmid for co-expression of GroEL and GroES | Takara Bio Inc. |
| pWTW009 | Deletion of *P. putida fadD1D2* (PP_4549-4550) | Linearized pk18mobsacB plasmid (oWWTW0010 & oWTW0011) assembled with homology arms from upstream (oWTW0038 & oWTW0039) and downstream (oWTW0036 & oWTW0037) of *fadD1D2* using Gibson assembly. |
| pWTW012 | Complementation of *P. putida fadD1* (PP_4549) | Bxb1 serine integrase cargo plasmid containing PP_4549 (*fadD*1) with 301 bp of the upstream region amplified from KT2440 gDNA (oWTW0113 & oWTW0115) and inserted into PCR linearized pWTW002 (oWTW0110 & oWTW0114) using Gibson assembly |
| pJH299 | Deletion of *P. putida alkK* (PP_0763) | EcoRI and HindIII linearized pk18mobsacB assembled with homology arms from upstream (oJH0722 & oJH0723) and downstream (oJH0724 & oJH0725) of *alkK* using Gibson assembly. |
| pJE365 | Deletion of *P. putida gcd* (PP_1444) | Elmore *et al.*, 2021[1] |

| | | |
|---|---|---|
| pJE1369 | Deletion of *P. putida hexR* (PP_1021) | EcoRI and HindIII linearized pJE382 (pk18mobsacB derivative) assembled with 700 bp homology arms from upstream (oJE1361 & oJE1362) and downstream (oJE1363 & oJE1364) of *hexR* using Gibson assembly. |
| pJH316 | Deletion of *P. putida phaG* (PP_1408) | Homology arms from upstream and downstream of *phaG* assembled into pk18mobsacB. DNA synthesis and cloning performed by Genscript. |
| pJE473 | Deletion of *P. putida phaC1ZC2* | Elmore *et al,*. 2021[1] |

**Table S3. DNA oligo sequences used in this study.**

Integrated DNA Technologies and Eurofins were used for synthesis.

| Primer | Sequence (5’- 3’) |
|---|---|
| oKM278 | ACCATCATCACCACAGCCAGAGCACTATCGAAGAACGCGT |
| oKM279 | AGGCGCGCCGAGCTCGTTACGCCTGGTGGCCGTT |
| oKM280 | ATAAGAAGGAGATATACATATGGCAATATTAGGTTTAGGCACGGA |
| oKM281 | GCGGTTTCTTTACCAGACTTAACTTTCAATAATTACCGTGGCACAAGC |
| oEUK116 | GCCGCGCGGCAGCCATATGACGCAATTTGCATTTG |
| oEUK117 | GTTAGCAGCCGGATCCTTAAAGCTCGAGCGCC |
| oEUK128 | GCCGCGCGGCAGCCATATGCATAACGTCGTGATCA |
| oEUK129 | GTTAGCAGCCGGATCCTCAGCGCTTGCGC |
| oEUK172 | AGAAGGAGATATACATATGAGCCTGCAAGGTAAAGTT |
| oEUK173 | AGCTCGAATTCGGATCCATGTACATCCCGCCGTT |
| oEUK174 | AGAAGGAGATATACATATGAGGCCAGAAATCGCTG |
| oEUK175 | AGCTCGAATTCGGATCCCAGATGGCAAATGCATGCTG |
| T7_fwd | TAATACGACTCACTATAGGG |
| T7_rev | GCTAGTTATTGCTCAGCGG |
| oWTW0010 | TGGCACTGGCCGTCGTTT |
| oWTW0011 | CGTAATCATGTCATAGCTGTTTCCTGAG |
| oWTW0036 | ACAGCTATGACATGATTACGCCATGGGCGGTATCGTCAC |
| oWTW0037 | ACAATAATAACCACAAGCCTGCACGCTC |
| oWTW0038 | AGGCTTGTGGTTATTATTGTTCCTCTGCCTGGACCTGC |
| oWTW0039 | TAAAACGACGGCCAGTGCCACGGTACAGCACGGCCGGT |
| oWTW0110 | GATCGCCTGAAACGCATGAGAAAGCCCC |
| oWTW0113 | CTCATGCGTTTCAGGCGATCTTCTTCAAG |
| oWTW0114 | GTGTAGGAGCCGGACCAAAACGAAAAAAGG |
| oWTW0115 | TTTTGGTCCGGCTCCTACACTGCCCAATG |
| oJH0722 | TCACTCAGGAAACAGCTATGACATGATTACGCATACCGAAGGCTTCGGCCAGCC<br>T |
| oJH0723 | GGATCCGTAACGTACTCTAGAACGGCCAAATCAGCCGAA |
| oJH0724 | TCTAGAGTACGTTACGGATCCTAACCATTGTGCGGGTCGCGC |
| oJH0725 | GTCACGACGTTGTAAAACGACGGCCAGTGCCACGACTTGGCGCCTTCCTT |
| (oJE1361)_pJE<br>1369_UP_F | CAGGAAACAGCTATGACATGATTACGAATTCTGTGGTGCGCCGAACA |
| (oJE1362)_pJE<br>1369_UP_R | GGACACACCCATGGACCGCGTGTGATTACCCGATCGAGGACGA |
| (oJE1363)_pJE<br>1369_DN_F | GTCGTCCTCGATCGGGTAATCACACGCGGTCCATGGGTGTGT |
| (oJE1364)_pJE<br>1369_DN_R | CGTTGTAAAACGACGGCCAGTGCCAAGCTTCGCCACGGCGTCGTT |

**Table S4. Strains and construction details for bacterial strains used in this study.**

| Strain | Genotype | Construction details |
|---|---|---|
| EUK148 | *E. coli* BL-21 λ(DE3) | *E. coli* BL-21 λ(DE3) transformed with pKM181 to produce *Ec*AcpP |
| EUK187 | *E. coli* BL-21 λ(DE3) | *E. coli* BL-21 λ(DE3) transformed with pEUK070 to produce *Ec*FabD |
| EUK201 | *E. coli* BL-21 λ(DE3) | *E. coli* BL-21 λ(DE3) transformed with pEUK079 to produce *Pp*FabH2 |
| EUK222 | *E. coli* BL-21 λ(DE3) | *E. coli* BL-21 λ(DE3) transformed with pEUK087 to produce *Pp*FabG |
| EUK225 | *E. coli* BL-21 λ(DE3) | *E. coli* BL-21 λ(DE3) transformed with pEUK086 & pGro7 to produce *Pp*PhaG |
| AG5577 | *P. putida* KT2440 ΔPP_2876::R4_phiBT1_MR11_attB cassette ΔPP_4740::Bxb1_RV_phi370_attB cassette ΔPP_4217/4218 intergenic::TG1_BL3_A118_attB cassette | Huenemann *et al.,* 2025[2] |
| AG6700 | *P. putida* KT2440 ΔPP_2876::R4_phiBT1_MR11_attB cassette ΔPP_4740::Bxb1_RV_phi370_attB cassette ΔPP_4217/4218 intergenic::TG1_BL3_A118_attB cassette Δ*alkK* (PP_0763) | *P. putida* AG5577 transformed with pJH299 (Δ*alkK,* PP_0763). Kanamycin was used to select for chromosomal integration, followed by sucrose counter-selection to select for the deletion. |
| AG7302 | *P. putida* KT2440 ΔPP_2876::R4_phiBT1_MR11_attB cassette ΔPP_4740::Bxb1_RV_phi370_attB cassette ΔPP_4217/4218 intergenic::TG1_BL3_A118_attB cassette Δ*fadD1D2* | *P. putida* AG5577 transformed with pWTW009 (Δ*fadD1D2*, PP_4549-4550). Kanamycin was used to select for chromosomal integration, followed by sucrose counter-selection to select for the deletion. |
| AG7303 | *P. putida* KT2440 ΔPP_2876::R4_phiBT1_MR11_attB cassette ΔPP_4740::Bxb1_RV_phi370_attB cassette ΔPP_4217/4218 intergenic::TG1_BL3_A118_attB cassette Δ*alkK* Δ*fadD1D2* | *P. putida* AG6700 transformed with pWTW009 (Δ*fadD1D2*, PP_4549-4550). Kanamycin was used to select for chromosomal integration, followed by sucrose counter-selection to select for the deletion. |
| AG7408 | *P. putida* KT2440 ΔPP_2876::R4_phiBT1_MR11_attB cassette ΔPP_4740::Bxb1_RV_phi370_attB cassette ΔPP_4217/4218 intergenic::TG1_BL3_A118_attB cassette Δ*phaC1ZC2* | *P. putida* AG5577 transformed with pJE473 (Δ*phaC1ZC2,* PP_5003-5). Kanamycin was used to select for chromosomal integration, followed by sucrose counter-selection to select for the deletion. |
| AG7670 | *P. putida* KT2440 ΔPP_2876::R4_phiBT1_MR11_attB cassette ΔPP_4740::Bxb1 attL-_RV_phi370_attB cassette ΔPP_4217/4218 intergenic::TG1_BL3_A118_attB cassette Δ*phaG* | *P. putida* AG5577 transformed with pJH316 (Δ*phaG,* PP_1408). Kanamycin was used to select for chromosomal integration, followed by sucrose counter-selection to select for the deletion. |
| AG8350 | *P. putida* KT2440 ΔPP_2876::R4_phiBT1_MR11_attB cassette ΔPP_4740::Bxb1attL-*fadD1*-attR_RV_phi370_attB cassette ΔPP_4217/4218 intergenic::TG1_BL3_A118_attB cassette Δ*alkK* Δ*fadD1D2* | *P. putida* AG7303 transformed with pWTW012 (Δ*fadD1,* PP_4549) |
| AG9017 | *P. putida* KT2440 ΔPP_2876::R4_phiBT1_MR11_attB cassette | *P. putida* AG7303 transformed with pΔphaZ (Δ*phaZ,* PP_5004) |

| | | |
|---|---|---|
| | ΔPP_4740::Bxb1_RV_phi370_attB cassette ΔPP_4217/4218 intergenic::TG1_BL3_A118_attB cassette Δ*alkK* Δ*fadD1D2* Δ*phaZ* | |
| AG10129 | *P. putida* KT2440 ΔPP_2876::R4_phiBT1_MR11_attB cassette ΔPP_4740::Bxb1_RV_phi370_attB cassette ΔPP_4217/4218 intergenic::TG1_BL3_A118_attB cassette Δ*alkK* Δ*fadD1D2* Δ*hexR* Δ*gcd* | *P. putida* AG7303 transformed with pJE1369 (Δ*hexR,* PP_1021) and pJE365 (Δ*gcd,* PP_1444) |

**Table S5. MRM transition table**

| Analyte name | Precursor ion | Ion | MRM quantifying transition (*m/z*) | Collision energy (V) | Fragmentor (V) | MRM qualifying transition (*m/z*) | Collision energy (V) |
|---|---|---|---|---|---|---|---|
| **3-Hydroxyhexanoic acid (C6-HA)** | 131.1 | [M-H]- | 131.1 → 59 | 8 | 80 | 131.1 → 129.9 | 0 |
| **3-Hydroxyoctanoic acid (C8-HA)** | 159.1 | [M-H]- | 159.1 → 59 | 12 | 95 | 159.1 → 157.8 | 4 |
| **3-Hydroxydecanoic acid (C10-HA)** | 187.1 | [M-H]- | 187.1 → 59 | 12 | 105 | 187.1 → 141 | 12 |
| **3-Hydroxydodecanoic acid (C12-HA)** | 215.2 | [M-H]- | 215.2 → 59 | 12 | 115 | 215.2 → 213.9 | 0 |
